# A Novel Method for the Mechanical Testing of Human Cerebrovascular Tissue: A Validation Study

**DOI:** 10.1101/2025.09.28.679090

**Authors:** Mark A Davison, Maximos P McCune, Nishanth Thiyagarajah, Daniel E Thomeer, Keyvon Rashidi, Majid Rashidi, Nina Z Moore

## Abstract

This study describes and validates a novel mechanical testing apparatus capable of generating a viscoelastic response from which the time-dependent behavior of human cerebrovascular tissue can be derived to inform vessel wall failure prediction and therapeutic device design. Testing involved vascular specimen cannulation, pressurization, and recording dynamic changes in vessel diameter using a three-axis laser micrometer. Device validity and versatility were evaluated via two synthetic microvessel specimen experiments: (I) comparison to an Instron stress-relaxation protocol, and (II) vessel segment length parametric analysis. A standard linear solid (SLS) model was chosen to fit the experimental results, from which the model coefficients (E_1_, E_2_, and *η*) and equilibrium modulus (G_e_) were computed. G_e_ comparisons were made using Bland-Altman analysis and Welch’s F-test for experiment I and II, respectively. Device feasibility was evaluated through testing human cadaveric cerebrovascular tissue. The SLS model provided accurate experimental data fits, with overall mean R^2^ value of 0.99 (SD= 2.4E-3). G_e_ for inflation-creep and Instron stress-relaxation experiments were statistically comparable, with Bland-Altman mean bias of 1.9% (95% CI: −0.9% - 4.6%, p=.18). Holistically, the vessel segment length parametric analysis revealed inconsistent values for G_e_ across the complete range of testing lengths, where ad hoc family-wise comparison indicated that the 0.5 cm length cohort was the singular outlier (p < .05). Our device successfully recorded a viscoelastic response from human cadaveric middle cerebral artery tissue (n=12). This study demonstrated that our novel device was both versatile and capable of eliciting an accurate viscoelastic response.

## INTRODUCTION

Ischemic stroke and intracranial hemorrhage are amongst the most prevalent causes of death and disability worldwide, the latter of which may be caused by intracranial vascular malformations such as aneurysms or arteriovenous malformations (AVMs) [1]. Contemporary treatments for intracranial vascular pathologies often consist of catheter based endovascular therapies which involve the deployment of a device that transiently or permanently alters the local vascular biomechanics and fluid dynamics to achieve a cure. As such, cerebrovascular biomechanics is quickly becoming a consequential field of study, as its implications influence novel medical device design and provide greater insight towards the underlying mechanisms responsible for the formation and stability of complex intracranial vascular pathologies [2–5].

While cerebrovascular tissue has been extensively characterized as nonlinear, viscoelastic and anisotropic, a more detailed understanding of the mechanical properties, particularly the time-dependent viscous behavior, is essential to enable continued advancement [6–7]. Preliminary work in cerebrovascular biomechanics has relied predominately on uniaxial tensile testing, which involves sectioning the vessel into planar strips and applying hooks or clamps to grasp the tissue, which tends to damage the specimen and create localized stress concentrations [8–12]. Most importantly, uniaxial experiments are limited in their capacity to capture the properties of a specimen that are known to be directional-dependent. As an alternative, inflation testing of a vessel segment facilitates a more physiologic, biaxial loading environment, which does not necessitate specimen disruption.

Multiple prior studies have successfully derived material properties of large extracranial and smaller intracranial animal vessels through inflation experiments [13–17]. Monson et al. were the first to translate their inflation testing methodology to freshly resected cerebral cortical vessels from human patients, however the study focused only on the axial and circumferential elastic response to pressurization [18]. Consequently, a comprehensive characterization of the viscoelastic properties of human cerebrovascular tissue, which would enable a more complete understanding of vessel hysteresis behavior under cyclic loading conditions, has yet to be established.

Unlike extracranial vascular tissue, the testing of cerebrovasculature represents a distinct challenge as it is exceptionally delicate, varies considerably in length, diameter and tortuosity, and is frequently associated with tiny perforating vessels easily avulsed despite meticulous dissection technique. Each of these factors has the tendency to create unavoidable mechanical testing constraints. Consequently, a device capable of producing accurate results while also maintaining the versatility to accommodate extensive specimen variation without introducing systematic inaccuracies is paramount. To that end, the purpose of this report was to propose and validate a novel mechanical testing apparatus capable of generating a viscoelastic profile from which the time-dependent behavior of human cerebrovascular tissue could be derived. Our concept involves creep experimental conditions with instantaneous vessel inflation serving as the stepwise load application.

## METHODS

### Mechanical Testing Device

The novel testing device utilized for these experiments involved instantaneously pressurizing a vessel segment with distilled water from a small reservoir and recording the change in vessel diameter at a cross-sectional plane of interest. Specifically, a vessel segment was mounted on a stainless-steel cannula and fixed with nylon suture at each end to prevent fluid leak. Distance between suture fixation points defined the effective specimen testing length and if desired, suture points could be shifted apart along the length of the cannula to introduce an axial pre-strain configuration (**Fig. 1**). The device featured interchangeable cannulas available in multiple diameters (range: 1 mm – 5 mm), which were selected according to the diameter of the vessel being tested. Each cannula contained boreholes about which the vessel segment was centered, which allowed pressurized fluid to enter the vessel lumen. Fluid from a pressure reservoir flowed into the proximal end of the cannula upon actuation of a solenoid valve, while the distal cannula end was fitted to a three-way valve, allowing the user to manually close or depressurize the system. The hydrostatic pressure within the vessel specimen was recorded by a 10 PSI pressure sensor (Walfront, Wuhan City, China) sampling at 10 Hz just proximal to the stainless-steel cannula. The vessel’s response to pressurization, an increase in diameter, was recorded using a three-axis non-contact laser micrometer device (LaserLinc, Fairborn, OH), which documented diameter measurements across three unique axes within a cross-sectional plane (**Fig. 2**). The timing of the laser micrometer measurements was synchronized to the 10 Hz pressure readings and diameters were averaged to compute a mean diameter measurement along the plane of interest, which was used to calculate instantaneous specimen strain. The experimental protocol was equivalent to a creep test, whereby the vessel was subjected to an instantaneous stepwise pressurization which was maintained until the average diameter reached a steady state, approximately 15 seconds after inflation. Between each inflation cycle, a cotton swab saturated in distilled water was applied to the vessel surface to ensure continuous vessel hydration.

**Figure 1.**
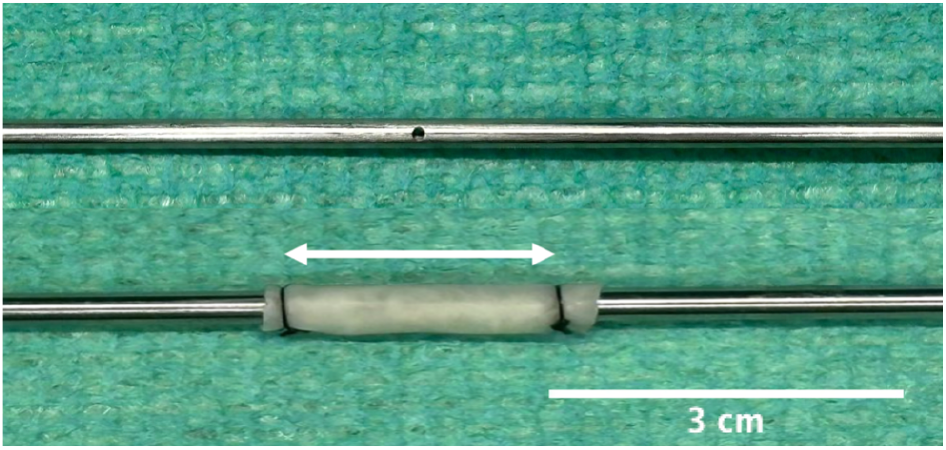
Stainless-steel cannula with borehole (top) and synthetic vessel fixed to cannula with nylon suture and double arrowheads demonstrating the effective specimen testing length (bottom).

**Figure 2.**
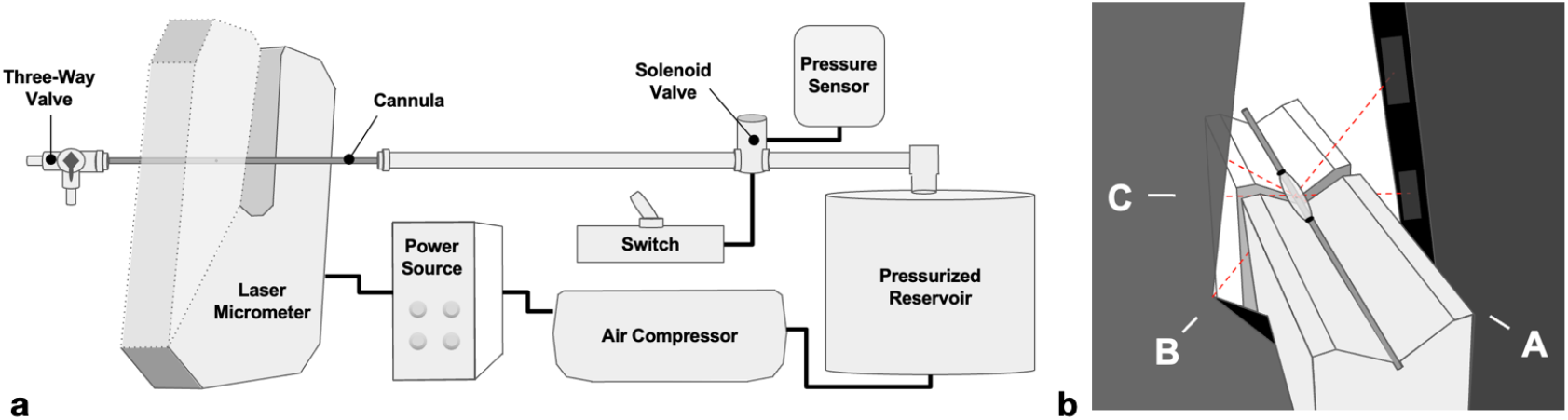
Schematic representation of the mechanical testing device (a); diagram demonstrating vessel mounted on cannula positioned within the laser micrometer (b).

### Synthetic Hydrogel Microvessel

The controlled experiments utilized a synthetic hydrogel microvessel engineered to resemble porcine vascular tissue (LifeLike BioTissue, London ON, CA), which was available in various outer diameters and wall thicknesses with consistent intrinsic properties. While the tissue grossly behaves as a non-linear viscoelastic and isotropic material, it exhibits a more linear behavior at lower strain magnitudes such as the ones employed in the current experiments. This material served as an ideal control for device validation as it was carefully manufactured to be a uniformly cylindrical without perforators or branching anatomy and was devoid of intimal atherosclerosis or other pathologic anomalies which may induce non-uniform biomechanical alterations. Synthetic vessel specimens were initially 7.6 cm long and ranged from 2 mm to 5 mm in outer diameter. Complete cross-sectional dimension specifications are provided in Appendix 1. Three pressurization cycles were sufficient to precondition the specimen and all inflation tests were performed without axial pre-strain, as the synthetic vessel naturally exists in a relaxed configuration.

### Constitutive Model Selection

The selection of a suitable constitutive equation was necessary to computationally model the synthetic vessel’s viscoelastic behavior and facilitate comparison across testing conditions. In choosing an experimental viscoelastic model, simplicity was paramount, as more complex rheological equations are computationally challenging and less conducive to translational research. While the Kelvin-Voight model represents the most fundamental viscoelastic solid, its ability to precisely capture the instantaneous elastic response following stepwise pressurization is limited. Alternatively, the Standard Linear Solid (SLS) model represented by one dashpot (*η*) and two spring elements (E_1_ and E_2_), serves as an improvement on the Kelvin-Voight model with the addition of a serial spring element to better capture the response to instantaneous loading (**Fig. 3**). The mathematical form of the SLS model is provided in Eq. 1.

**Figure 3.**
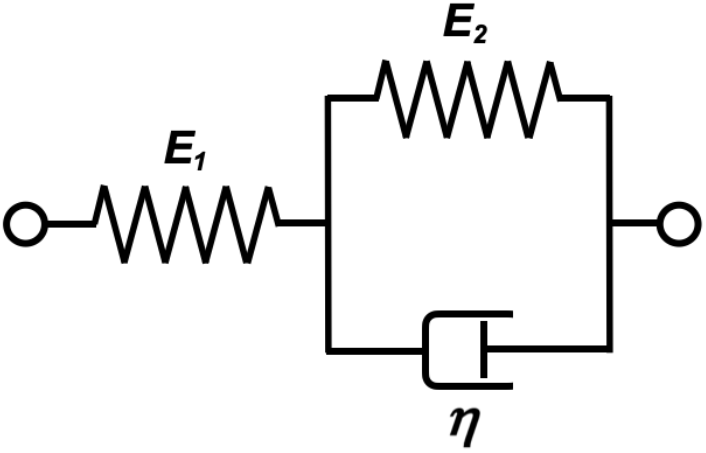
Schematic representation of the Standard Linear Solid model, with spring (E_1_ and E_2_) and dashpot (η) elements.

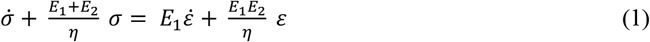

When subjecting a viscoelastic material, such as the synthetic hydrogel microvessels, to a sustained loading condition, the time-dependent viscous portion of its properties gradually settle and the material’s effective stiffness approaches a steady-state. This effective stiffness is represented by the equilibrium modulus (G_e_), which takes the form of Eq. 2 for the SLS model described above.

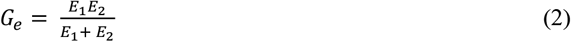

Prior to the current experiments, the equilibrium modulus was selected as the material property by which subgroup comparisons would be made. The rationale for this decision was as follows: G_e_ represented a single metric and therefore results were unequivocal. Secondly, G_e_ could be measured with a high degree of confidence as the microvessel tissue response observed during pilot trials resemble a step-function, suggesting that the SLS element η would likely be insensitive and relatively inconsistent.

### Device Validation Overview

To thoroughly assess the validity and versatility of our novel device, two controlled experiments were performed using a synthetic microvessel specimen with the following objectives: (I) demonstrate that the equilibrium modulus was comparable to the values obtained from a standard stress-relaxation testing experiment, and (II) prove that the calculated equilibrium modulus values were consistent and independent of the effective vessel testing length. As a final test to verify device feasibility, a series of human cadaveric cerebrovascular tissue samples were subjected to our inflation-creep protocol in an attempt to elicit a viscoelastic response.

### Controlled Validation Experiments

#### I. Equilibrium Modulus Comparison

Verifying the ability of the device to derive accurate material properties from an experimental viscoelastic response was a fundamental validation step. To accomplish this, results derived from our inflation-creep experiments would be compared to a matched series of uniaxial stress-relaxation tests performed using an Instron 68TM-30 with 2530 static 30 kN load cell (Instron, Noorwood, MA). In this case, as the synthetic microvessels behave as an isotropic material, it was appropriate to utilize uniaxial testing conditions as the gold standard experimental methodology. Matched specimen pairs were established by transecting a synthetic vessel of 7.6 cm length, with one part allocated for an inflation-creep test and the other larger segment designated for an Instron stress-relaxation experiment. Testing was performed on three, 5 mm outer diameter vessels which were best suited for the Instron clamps (specimens I.A–I.C, **Table 1**).

**Table 1.** Testing protocol for each synthetic microvessel validation experiment: Equilibrium Modulus Comparison (I.A-I.C) and Vessel Segment Length Parametric Analysis (II.A-II.C).

| Specimen ID | Testing Configuration | Outer Diameter (mm) | Effective Testing Length (cm) | Inflation Pressure (PSI) | Testing Cycles (n) |
| --- | --- | --- | --- | --- | --- |
| Equilibrium Modulus Comparison |  |  |  |  |  |
| I.A | Inflation Creep Test | 5 | 1.0 | 2.5 | 10 |
| I.B | Inflation Creep Test | 5 | 1.0 | 3.5 | 10 |
| I.C | Inflation Creep Test | 5 | 1.0 | 4.5 | 10 |
| I.A | Instron Stress-Relaxation Test | 5 | 4.5 | * N/A | 10 |
| I.B | Instron Stress-Relaxation Test | 5 | 4.5 | † N/A | 10 |
| I.C | Instron Stress-Relaxation Test | 5 | 4.5 | ‡ N/A | 10 |
| Vessel Segment Length Parametric Analysis |  |  |  |  |  |
| II.A | Inflation Creep Test | 4 | 2.5 | 2.5 | 10 |
| II.A | Inflation Creep Test | 4 | 2.0 | 2.5 | 10 |
| II.A | Inflation Creep Test | 4 | 1.5 | 2.5 | 10 |
| II.A | Inflation Creep Test | 4 | 1.0 | 2.5 | 10 |
| II.A | Inflation Creep Test | 4 | 0.75 | 2.5 | 10 |
| II.A | Inflation Creep Test | 4 | 0.5 | 2.5 | 10 |
| II.B | Inflation Creep Test | 4 | 2.5 | 2.5 | 10 |
| II.B | Inflation Creep Test | 4 | 2.0 | 2.5 | 10 |
| II.B | Inflation Creep Test | 4 | 1.5 | 2.5 | 10 |
| II.B | Inflation Creep Test | 4 | 1.0 | 2.5 | 10 |
| II.B | Inflation Creep Test | 4 | 0.75 | 2.5 | 10 |
| II.B | Inflation Creep Test | 4 | 0.5 | 2.5 | 10 |
| II.C | Inflation Creep Test | 4 | 2.5 | 2.5 | 10 |
| II.C | Inflation Creep Test | 4 | 2.0 | 2.5 | 10 |
| II.C | Inflation Creep Test | 4 | 1.5 | 2.5 | 10 |
| II.C | Inflation Creep Test | 4 | 1.0 | 2.5 | 10 |
| II.C | Inflation Creep Test | 4 | 0.75 | 2.5 | 10 |
| II.C | Inflation Creep Test | 4 | 0.5 | 2.5 | 10 |

Stepwise Instron displacements for stress-relaxation experiments: \*1.0 cm (22% strain), †1.5 cm (33% strain), and ‡ 2.0 cm (44% strain).

**Table 2.**
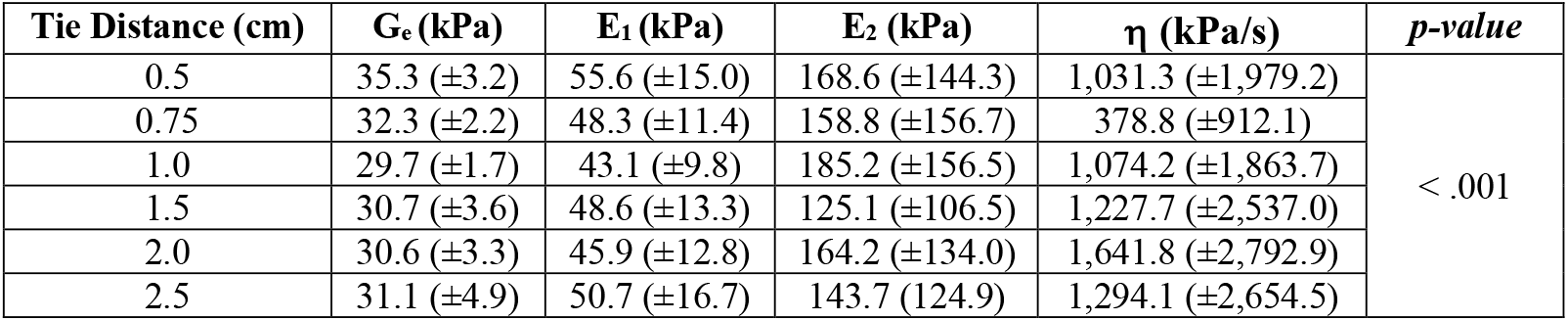
Calculated equilibrium modulus and SLS coefficients from vessel segment length parametric analysis experiments, where p-value was determined via Welch’s F-test.

| **Tie Distance (cm)** | **G<sub>e</sub> (kPa)** | **E<sub>1</sub> (kPa)** | **E<sub>2</sub> (kPa)** | **η (kPa/s)** | ***p-value*** |
| --- | --- | --- | --- | --- | --- |
| 0.5 | 35.3 (±3.2) | 55.6 (±15.0) | 168.6 (±144.3) | 1,031.3 (±1,979.2) | < .001 |
| 0.75 | 32.3 (±2.2) | 48.3 (±11.4) | 158.8 (±156.7) | 378.8 (±912.1) |  |
| 1.0 | 29.7 (±1.7) | 43.1 (±9.8) | 185.2 (±156.5) | 1,074.2 (±1,863.7) |  |
| 1.5 | 30.7 (±3.6) | 48.6 (±13.3) | 125.1 (±106.5) | 1,227.7 (±2,537.0) |  |
| 2.0 | 30.6 (±3.3) | 45.9 (±12.8) | 164.2 (±134.0) | 1,641.8 (±2,792.9) |  |
| 2.5 | 31.1 (±4.9) | 50.7 (±16.7) | 143.7 (124.9) | 1,294.1 (±2,654.5) |  |

Inflation-creep tests were completed first, where a 5 mm diameter synthetic vessel was cannulated and secured with nylon ties, forming an effective testing length of 1.0 cm. Once preconditioned, the specimen (I.A) was pressurized to 2.5 PSI and maintained until a steady state mean diameter was achieved. This experiment was repeated for a total of ten testing cycles. Two additional 5 mm diameter specimens were then tested at 3.5 PSI (I.B) and 4.5 PSI (I.C), likewise with a 1.0 cm effective testing length. Next, Instron stress-relaxation experiments were performed. A 5 mm diameter specimen was carefully secured in no-slip smooth clamps with an initial relaxation length of 4.5 cm. Following adequate preconditioning, specimen I.A was subjected to an instantaneous stepwise displacement of 1.0 cm (22.2% strain) and the resultant load response was continuously recorded for 15 seconds. The specimen was returned to its relaxation length and the testing cycle was repeated for a total of ten cycles. This process was replicated for the remaining two synthetic specimens, with stepwise displacements of 1.5 cm (33.3% strain) and 2.0 cm (44.4% strain), which corresponded to specimen I.B and I.C, respectively (**Table 1**). These strain magnitudes were selected to resemble the experimental strain observed in the inflation-creep experiments.

The equilibrium modulus (G_e_) was calculated from the inflation-creep and stress-relaxation experimental viscoelastic profiles, matched by specimen and cycle number, and compared graphically. Additionally, matched experimental data was compared using Bland-Altman analysis, a well-established method for assessing the degree of agreement between two quantitative measurement protocols [19]. The mean percent difference for each modulus pair was calculated and assessed for a fixed measurement bias via a single sample t-test, with a value of 0 indicating no difference. This test served as the limits of agreement criteria.

#### II. Vessel Segment Length Parametric Analysis

To verify our device reproduced consistent results irrespective of the effective specimen testing lengths, systematic inflation tests were conducted. Using a cannulated 4 mm outer diameter synthetic vessel, nylon ties were first placed to create a 2.5 cm effective testing length. Once preconditioned, ten testing cycles were completed at 2.5 PSI. Similarly, subsequent tests were performed for effective testing lengths of 2.0 cm, 1.5 cm, 1.0 cm, 0.75 cm, and 0.5 cm, and this protocol was repeated on a total of three synthetic specimens (II.A-II.C, **Table 1**). Likewise, the equilibrium modulus was calculated for each loading cycle and analyzed with respect to effective testing length.

### Human Cerebrovascular Tissue Experiment

Cerebrovascular tissue was obtained from a saline-flushed fresh-frozen cadaver cephalad specimen (Science Care, Phoenix, AZ). Following careful dissection and procurement of tissue from the proximal and distal regions of the bilateral middle cerebral artery circulation, the specimens were cryopreserved in dimethylsulfoxide (DMSO) and stored in a −80°C freezer. Prior to mechanical testing, samples were slowly thawed to 37°C in a water bath. The specimen was examined for a uniform segment free of tiny perforating or avulsed branches, carefully mounted onto an appropriately sized cannula and fastened at each end with nylon suture to create an effective specimen length between 1.0 and 2.0 cm. The vessel was inflated to 2.5 PSI and its diameter was recorded until a steady state was observed. This process was repeated for each of the dissected middle cerebral artery specimen (n = 12).

### Data Processing and Statistical Analysis

Experimental measurements of time, diameter and pressure were imported into MATLAB version R2023a (MathWorks, Natick, Massachusetts), where experimental curves were appropriately trimmed, and circumferential stress and strain were calculated from the mean cross section diameter and hydrostatic pressure changes. Processed data was curve fit to the three SLS equation coefficients (E_1_, E_2_, and η) using a minimization function based on the sum of squared errors. From these experimental coefficients, G_e_ was computed for each synthetic vessel test. A matched equilibrium modulus (G_e_) comparison between the inflation-creep and stress-relaxation experiments was conducted using a graphical approach and a pairwise Bland Altman analysis as described above. For the vessel segment length parametric analysis, box plots were generated depicting the mean, median, first and third quartiles, as well as maximum and minimum value ranges. Additionally, parametric cohort comparisons were made using Welch’s F-test under the assumption of unequal variances, with an ad hoc family-wise comparison if indicated. All statistical comparisons were two-sided and p < 0.05 was considered significant. Statistical calculations were performed using R version 4.3.1 (R Foundation, Vienna, Austria).

## RESULTS

The mechanical testing device was able to instantaneously inflate a synthetic microvessel specimen to a pre-defined hydrostatic pressure, maintain a constant pressurization for a desired length of time, and record a viscoelastic response via the laser micrometer (**Figure 4**). Notably, despite disparate specimen effective testing lengths, the experimental viscoelastic profiles remain consistent. Regarding the SLS model coefficients, the E_1_ term predominately captured the initial linear elastic response, while the secondary viscoelastic character was described by the interaction of the E_2_ and η terms. Generally speaking, during this time-dependent phase of the specimen’s behavior, the E_2_ term dictated the magnitude of strain the vessel experiences during the pressurization experiment, while the η coefficient modified the profile the strain response takes to get to this value.

**Figure 4.**
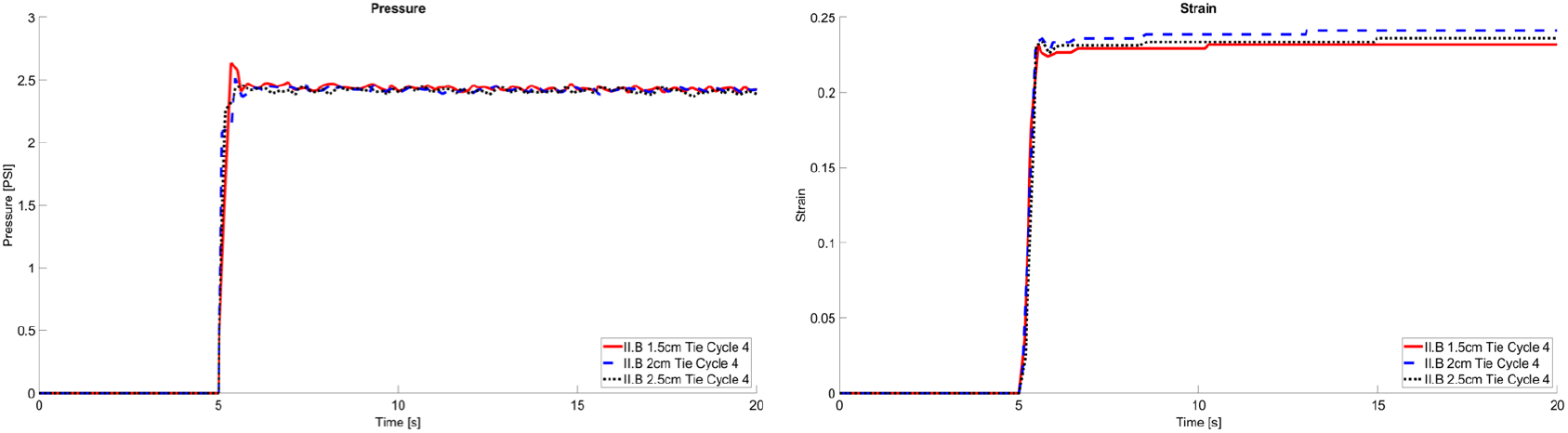
Pressure versus time (left) and strain versus time (right) for three representative synthetic vessel inflation-creep experiments (II.B: 1.5 cm tie, cycle 4; II.B: 2.0 cm tie, cycle 4; II.B: 2.5 cm tie cycle 4).

### Controlled Synthetic Microvessel Validation Experiments

#### I. Equilibrium Modulus Comparison

The SLS model provided an accurate fit for both the inflation-creep and stress-relaxation experiments, with mean R^2^ values of 0.99 (SD ± 3.9E-5) and 0.99 (SD ± 5.4E-4), respectively. The average inflation-creep testing E_1_, E_2_, η coefficient values were 46.3 kPa (SD ± 19.8), 137.4 kPa (SD ± 85.6), and 259.9 kPa-s (SD ± 102.9), respectively. Similarly for the stress-relaxation experiments, mean values for E_1_, E_2_, η were 39.9 kPa (SD ± 6.5), 66.0 kPa (SD ± 11.2), and 42.2 kPa-s (SD ± 43.9), respectively. Finally, mean equilibrium moduli for the inflation-creep and stress-relaxation experiments were 24.9 kPa (SD ± 2.4 kPa) and 24.5 kPa (SD ± 2.7 kPa), respectively. On comparison, the experimentally obtained equilibrium moduli values, G_e_, were comparable between the two methodologies when visualizing the results graphically and according to the Bland-Altman analysis. Specifically, the mean bias for G_e_ was 1.9% (95% CI: −0.9% - 4.6%, p = .18), indicating no statistical discrepancy between the two mechanical testing processes. Moreover, there were no discernible patterns or trends between strain magnitude and measurement disagreement (**Figure 5**).

**Figure 5.**
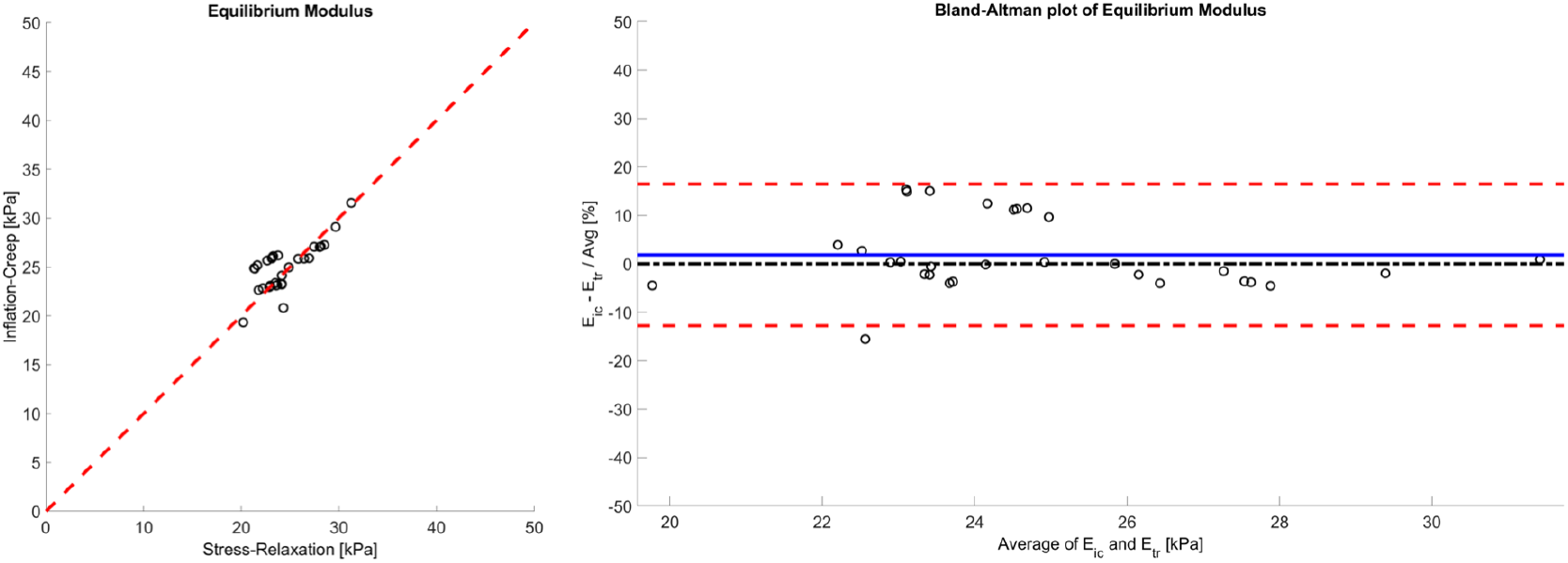
Graphical representation of equilibrium moduli derived from stress-relaxation experiments versus inflation-creep experiments with dashed y = x reference line serving as visual representation (left-hand side); Pairwise Bland Altman plots comparing equilibrium moduli from the two experimental methodologies, with solid line indicating the average percent difference and the dashed lines representing ± 1.96 * standard deviation limits of agreement (right-hand side).

#### II. Vessel Segment Length Parametric Analysis

SLS model fits for the included experiments were satisfactory, with a mean R^2^ value of 0.99 (SD ± 2.3E-5). Average model coefficients E_1_, E_2_, and η were 48.8 kPa (SD ± 14.0), 157.5 kPa (SD ± 136.2), and 1,171.6 kPa/s (SD ± 2,295.5), respectively, with an average experimental G_e_ of 31.6 kPa (SD ± 3.9). The mean experimental coefficients and equilibrium modulus for each effective vessel length testing condition are provided in **Table 3**. On overall cohort comparison, Welch’s F-test indicated that G_e_ was not consistent across all specimen testing lengths (p < .001). Subsequent family-wise comparisons revealed that the equilibrium modulus was statistically comparable for effective vessel lengths 2.5 cm through 0.75 cm, with vessels tested at length of 0.5 cm deviating from each of the other length groupings (p < .05), (**Figure 6**).

**Figure 6.**
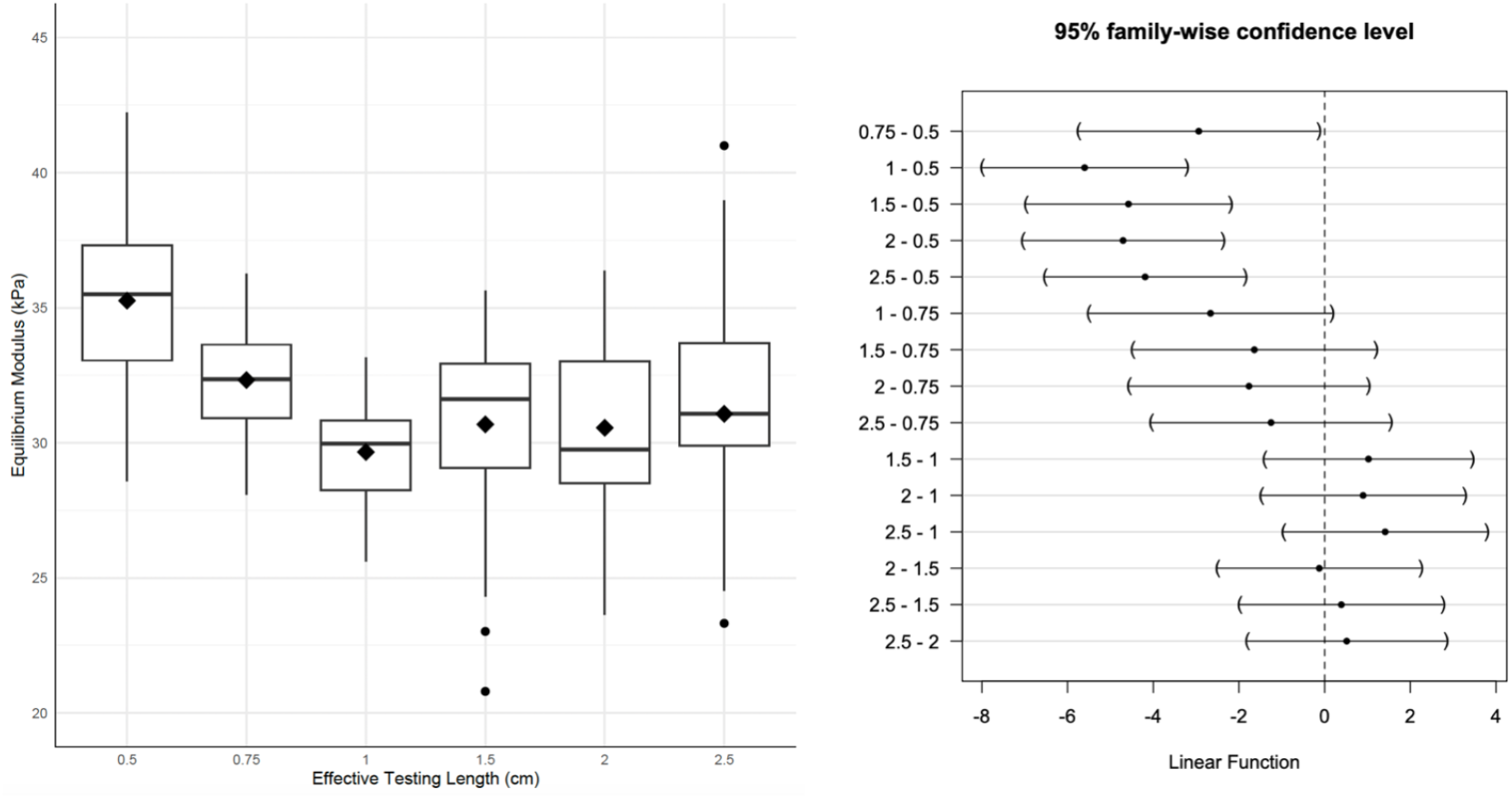
Vessel segment length parametric analysis box plot comparison for G_e_, where the mean and median are represented by the diamond and bold line, respectively and the whiskers indicate the range (left-hand side). Family-wise comparison for each vessel length cohort match-up based on 95% confidence intervals (right-hand side).

### Human Cerebrovascular Tissue Experiments

The inflation-creep testing methodology was successfully replicated in all 12 specimens derived from cadaveric cerebrovascular tissue from the middle cerebral artery distribution. More specifically, we were able to instantaneously inflate the cadaveric vessel segment and maintain a constant hydrostatic pressure for a desired amount of time, which exceeded 10 seconds in the current experiment. The laser micrometer acquired cross-sectional measurements from the vessel surface, enabling the recording of a viscoelastic response under creep testing conditions (**Figure 7, 8**). Moreover, the device and protocol were versatile enough to accommodate cerebrovascular tissue from throughout the middle cerebral artery vascular territory, which varied considerably in diameter, tortuosity, and morphology.

**Figure 7.**
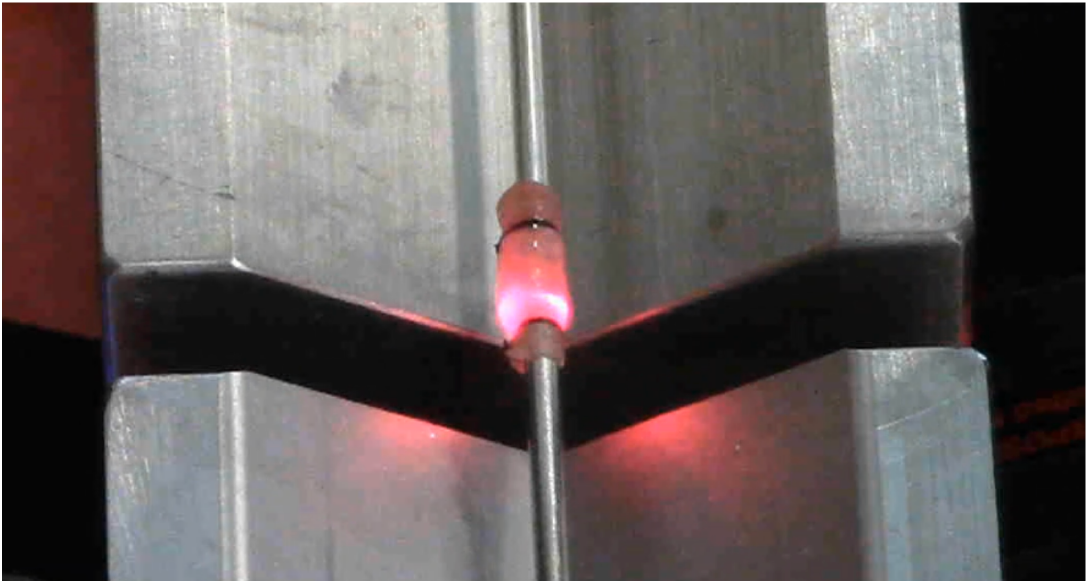
Image depicting an inflated human cadaveric vessel specimen from the middle cerebral artery circulation with laser micrometer recording three diameter measurements within a single cross-sectional plane of interest.

**Figure 8.**
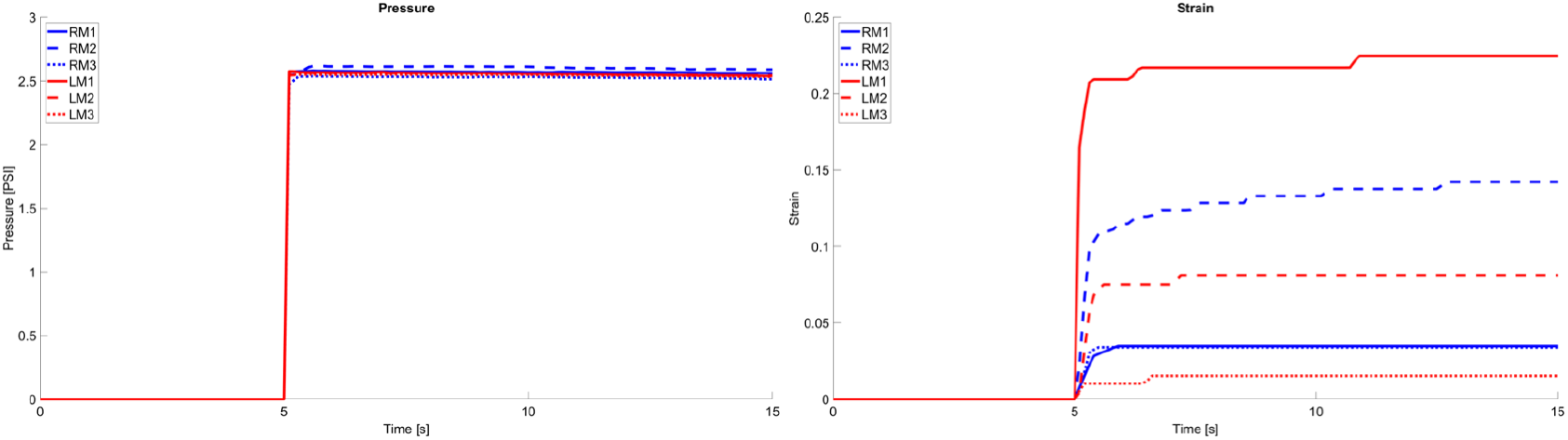
Pressure versus time (left) and strain versus time (right) from inflation-creep experiments of six representative cadaveric vessels from the left and right sided middle cerebral artery vascular territory, from proximal (M1), middle (M2), and distal (M3) segments.

## DISCUSSION

In this study, we proposed a novel mechanical testing device capable of generating a viscoelastic response under inflation-creep experimental conditions, utilizing both a synthetic microvesssel and human cadaveric cerebrovascular tissue. Moreover, we successfully validated the accuracy and versatility of our device through a rigorous series of controlled experiments. To our knowledge, this is the first device of its kind to experimentally derive viscoelastic properties from a microvessel specimen through an inflation testing protocol.

In the current experiment, we demonstrated that the true equilibrium modulus of a synthetic microvessel could be extracted from our inflation data through a matched analysis, using Instron stress-relaxation experiments as the standard. Specifically, calculated G_e_ values were highly consistent between protocols, with an average Bland-Altman bias of 1.9%, which was statistically negligible. Likewise, we established the adaptability of our testing apparatus through a vessel segment length parametric analysis, whereby the equilibrium modulus remained consistent for testing lengths between 0.75 cm and 2.5 cm. Interestingly, at effective specimen lengths of 0.5 cm, the mean G_e_ increased in relation to the 0.75 cm – 2.5 cm testing conditions, which would suggest a limiting physical constraint to the current experimental configuration. This may be related to the effective vessel geometry deviating from a cylindrical morphology and approaching a more spherical configuration, necessitating an alternative modelling approach.

While there is a long history of approximating the material properties of cerebrovascular tissue through mechanical testing experiments, the utility of these findings is limited by methodological deficiencies. As an anisotropic tissue, the stress-response behavior of cerebrovascular tissue cannot be sufficiently characterized through uniaxial loading conditions alone. Alternatively, biaxial testing experiments permit a more robust understanding of the complex directional-dependent material properties we know to exist. Previous studies have subjected planar sections of extracranial vascular tissue to biaxial loading, which often involve dual-axis tensile tests requiring the application of hooks or clamps which introduce local tissue damage and stress concentrations resulting in data artifact [16, 20–22]. Marra et al. proposed an alternative experimental protocol whereby a prepared planar section of vascular tissue is circumferentially fastened to a plastic washer and inflated in order to measure rupture strength [23]. While the latter method utilizes more uniform boundary conditions, each of these experiments necessitates disruption of the tissue’s innate anatomic structure and is therefore poorly suited for characterizing whole vessel biomechanics.

A review of the vascular mechanics literature revealed two studies investigating the biaxial response and elastic properties of freshly resected human cerebrovascular tissue through combined axial extension-inflation experiments [18, 24]. While these proposed methodologies for mechanical testing serve as a significant advancement, they are not without their own limitations. Monson et al. outlined an original, albeit extensive specimen preparation process that required the application of cyanoacrylate at the cannulated specimen ends, meticulous placement of 10 µm microspheres on the vessel surface, and careful suture ligation of delicate perforating branches [18]. While this protocol was suitable for their specific aims which centered around cortical vessel injury, these vessels tend to be smaller, more uniformly cylindrical and less tortuous.

The work of Liu et al. alternatively utilized a pressure myograph for their axial extension-inflation testing, a device initially designed to study the vasoactive behavior of small vessels under dynamic physiologic and pharmacologic environments. In these experiments, the authors studied the effects of amyloid-β deposition on the mechanical response of distal anterior cerebral arteries in patients with Alzheimer’s disease [24]. While pressure myography also permits the testing of an intact vascular specimen, the tissue preparation process necessitates suture ligation of the vessel ends onto two tiny glass cannulas, which are recessed in a 20 mL fluid chamber; an exacting task requiring an operating microscope, microsurgical instruments and advanced bimanual dexterity [25]. Moreover, pressure myography is specifically designed for assessing small diameter vessels and as such, is limited in its ability to assess the elastic and viscoelastic properties of the more proximal circle of Willis vasculature with thicker walls, intimal plaque, tortuous nature and an abundance of perforators. Consequently, despite these advancements in mechanical testing protocol, it was evident that an alternative process was required to satisfy our objectives.

The primary motivation for our device and protocol design was to create methodology capable of producing a viscoelastic response from human cerebrovascular tissue under inflation-creep loading conditions. Equally essential was ensuring the device was versatile enough to accommodate a diversity of specimens from throughout the intracranial circulation, could seamlessly adapt to physical testing constraints, and involved a simple and minimally destructive specimen preparation process that would facilitate efficient testing of a large quantity of cadaveric vessels and surgically resected pathology. Our novel method of tissue support, a single continuous cannula, enables the testing of an extensive range of human cerebrovascular tissue including thin-walled delicate vessels as well as specimens of sub-centimeter tissue lengths. Moreover, rather than pressurizing the specimen through cannulated ends, our design allows the vessel to be inflated through a central borehole, which mitigates the edge effects observed when fluid flows from a rigid cannula lumen into a biologic vessel and vice versa. The established protocol merely requires cannulating a specimen and ligating the two ends with nylon suture, which can be done on a table top and without specialized instrumentation. When desired, axial pre-strain can be introduced by simply sliding the specimen ligation points apart along the length of the cannula. The selection of laser optometry as the mechanism for documenting the tissue strain response is a novel advancement which obviates the need for microsphere application [13, 18] and provides a more objective measurement with greater resolution when compared to ultrasonography [14, 26]. Finally, the strain computation requires minimal data post-processing, further improving procedural efficiency.

While cerebrovascular biomechanics is a rapidly evolving field of study, there remains an immense potential for continued advancement and application of acquired knowledge towards areas of considerable clinical and scientific importance. For example, with a more comprehensive characterization of the time-dependent mechanical properties of human cerebrovasculature tissue, we may enhance our understanding of the origins and behavior of complex intracranial vascular pathologies. It has been demonstrated that the natural history and high-risk features of a vascular malformation may correlate with its precise intracranial location [27–28], an observation supported by fundamental concepts in cerebrovascular embryology, as different regions of the intracranial vasculature derive from disparate origins [29]. Computational fluid structure interaction (FSI) modelling, a method for simulating the complex interactions between a dynamic fluid and its structural surroundings, represents a promising tool for predicting the stability of such cerebrovascular pathologies [30–31]. While early applications of this technique have been successful [5], an improved understanding of the viscoelastic behavior of normal and pathologic cerebrovascular tissue through methods like the one described in this report will significantly enhance FSI modelling capabilities and ultimately improve clinical care [32].

## Limitations

The findings of this study must be interpreted within context of its assumptions and limitations. The inflation testing device records diameter measurements at a single cross-sectional plane within the mid-section of the specimen, which assumes the specimen is uniform and the strain is homogenous throughout the effective testing length. While this is more likely to be the case for a manufactured synthetic specimen, this is less likely true for a resected human cerebral vessel with length dependent mural inconsistencies. Reviewing the inflation-creep results, there were observed pressure losses throughout the duration of the testing cycles, which were proximal to the cannula and not related to fluid leak through the ligated ends of the specimen. While this may affect the recorded viscoelastic response curves, the magnitude of pressure loss was less than 5% and therefore deemed negligible. Finally, in similar experiments involving intact vascular specimen inflation, the tissue is submerged in a small fluid bath throughout the duration of the testing cycle. While it is possible the additional ambient, unpressurized fluid environment may influence our findings, it was felt the effects would be negligible and we prioritized obtaining frequent and accurate laser micrometer diameter measurements which necessitated testing outside of a liquid chamber.

## APPENDIX

**Appendix 1**. Cross-sectional dimensions of the synthetic microvessel specimens utilized in the validation experiments, obtained from manufacturer.

| Outer Diameter (mm) | Wall Thickness (mm) |
| --- | --- |
| 2.0 | 0.25 |
| 3.0 | 0.25 |
| 4.0 | 0.50 |
| 5.0 | 0.50 |

